# Template-free prediction of a new monotopic membrane protein fold and oligomeric assembly by Alphafold2

**DOI:** 10.1101/2022.07.12.499809

**Authors:** Alican Gulsevin, Bing Han, Jason C. Porta, Hassane S. Mchaourab, Jens Meiler, Anne K. Kenworthy

**Affiliations:** Department of Chemistry, Vanderbilt University Nashville, TN, USA; Center for Membrane and Cell Physiology, University of Virginia, Charlottesville, VA USA; Department of Molecular Physiology and Biological Physics, University of Virginia School of Medicine, Charlottesville, VA, USA; Life Sciences Institute, University of Michigan, Ann Arbor, MI, USA; Department of Molecular Physiology and Biophysics, Vanderbilt University Nashville, TN, USA; Institute for Drug Discovery, Leipzig University, Germany

## Abstract

AlphaFold2 (AF2) has revolutionized the field of protein structural prediction. Here, we test its ability to predict the tertiary and quaternary structure of a previously undescribed scaffold with new folds and unusual architecture, the monotopic membrane protein caveolin-1 (CAV1). CAV1 assembles into a disc-shaped oligomer composed of 11 symmetrically arranged protomers, each assuming an identical new fold, and contains the largest parallel β-barrel known to exist in nature. Remarkably, AF2 predicts both the fold of the protomers and interfaces between them. It also assembles between 7 and 15 copies of CAV1 into disc-shaped complexes. However, the predicted multimers are energetically strained, especially the parallel β-barrel. These findings highlight the ability of AF2 to correctly predict new protein folds and oligomeric assemblies at a granular level while missing some elements of higher order complexes, thus positing a new direction for the continued development of deep learning protein structure prediction approaches.

## Introduction

A long-standing goal of computational biology is to accurately predict the three-dimensional structure of proteins from their primary sequence. This goal appears now within reach using deep learning algorithms trained on a database of known proteins, most prominently AlphaFold2 (AF2) [1]. AF2 can predict protein structure even in the absence of high-homology templates [1, 2]. Another advantage of AF2 is its ability to predict multimeric assemblies without prior knowledge of the interaction sites of partner proteins or subunits [3, 4]. This feature is useful when the quaternary structure of a multimeric protein is unknown.

Despite the many successes of AF2 [5-8], its performance with structures for which there are no homologs present in the PDB remains an open question. Although AF2 does not rely on templates for structure prediction, the AF2 algorithm was trained based on available structural information from most structures in the PDB [1]. This may potentially limit its ability to model structures that fall outside the scaffolds of these proteins. One such example is structures obtained through NMR, which were not included in the training set of AF2. While AF2 was mostly successful in predicting these protein structures, it had lower accuracy in some cases, especially with flexible proteins [9]. In other cases, lower accuracy predictions by AF2 were argued to be related to a less deep MSA, high proportion of heterotypic contacts, and high oligomerization states [10]. Therefore, studies focusing on the prediction of novel protein scaffolds with AF2 can shed light on both its strengths and areas for future improvement.

We recently determined a cryo-EM structure for the protein caveolin-1 (CAV1), an integral membrane protein that plays a role in formation of membrane invaginations called caveolae [11]. CAV1 has several unusual and unexpected features in its structure that make it exceptionally well suited to benchmark the ability of AF2 to predict the structure of proteins with unusual scaffolds. First, CAV1 is a member of a protein family that shares little sequence or structural homology to other proteins [11], and no high resolution structures of any caveolin family members were available in the PDB at the time AF2 was trained. Second, CAV1 is a monotopic membrane protein, a class of proteins which are underrepresented in the PDB compared to other types of membrane proteins [12]. Third, the CAV1 complex has a previously undescribed quaternary structure, consisting of a tightly packed disc composed of 11 primarily α-helical protomers that tightly pack into a spiral arrangement. Finally, the complex contains an 11-fold symmetric parallel β-barrel, the largest parallel β-barrel to be reported to date. This distinguishes it from known examples of parallel β-barrel structures, belonging primarily to the TIM barrel family whose members consist of eight β-strands that fold into a barrel with four-fold symmetry [13].

In this work, we used CAV1 as a test case to address two questions: The first one is whether AF2 can model the tertiary and quaternary structure of a scaffold for which it has no homologs in its training set. The second question is whether the CAV1 models generated by AF2 can be used to understand the mechanism of CAV1 oligomerization.

## Results

### The unique experimental CAV1 structure presents a formidable test case for AF2

We previously determined the only experimental structure of the human CAV1 complex (PDB-7SC0). It consists of 11 CAV1 protomers, each of which assumes an identical new fold, that are symmetrically arranged into a tightly packed disc-shaped complex with a central pore formed by an 11-stranded parallel β-barrel [11] (**Figure 1A, B**). Residues 1 to 48 of the 178 residue protomers are absent from the cryo-EM structure and are predicted to be unstructured [14, 15]. Structurally and functionally important regions of the protein include the pin motif (PM, residues 49 to 60), a loop that envelopes and stabilizes the interactions between neighboring protomers; the caveolin signature motif (SM, residues 68 to 75), a highly conserved region across all caveolins composed of short α-helical region; and the scaffolding domain (SD, residues 82-101), comprising a portion of α-helix 1 at the periphery of the CAV1 8S disc that assists with oligomerization as part of a larger oligomerization domain (OD, residues 61-101). Other notable domains include a region traditionally referred to as the ‘intra-membrane’ domain (IMD, residues 102-134), consisting of a helical region that defines one boundary of the membrane facing surface of the protein, a long C-terminal amphipathic α-helix termed the spoke region (SR, residues 135-169), and a C-terminal β-strand (residues 170-176) that contributes to the assembly of the central β-barrel in the complex [11].

**Figure 1:**
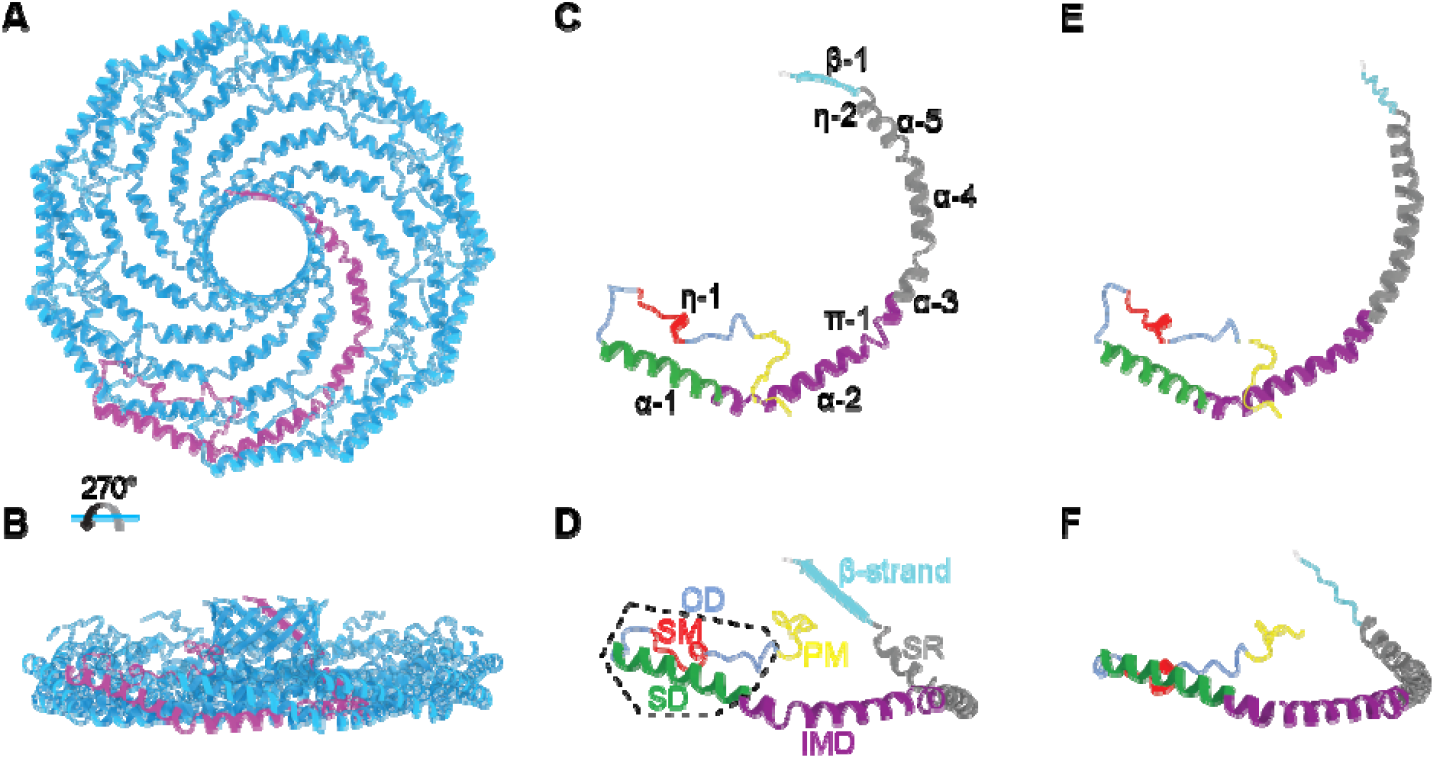
The structure of a CAV1 monomer predicted by AF2.1 closely resembles that of a CAV1 protomer experimentally determined by cryo-EM. (**A, B**) Top and side view of human CAV1 8S complex cryo-EM structure (PDB 7SC0) with a single protomer colored purple. (**C, D**) Top and side views of a single CAV1 protomer extracted from the experimental structure. The secondary structure assignments for the cryo-EM structure are labeled. Specific regions of CAV1 are labeled and colored as follows: PM, pin motif (yellow); SM, signature motif (red); SD, scaffolding domain (green); IMD, intermembrane domain (purple); SR, spoke region (grey); and β-strand (cyan). The OD, which contains the SM and SD, is indicated by the dashed box. (**E, F**) Top and side views of the AF2.1-predicted CAV1 monomer (AF-Q03135).

### AF2 predicts the tertiary structure of the CAV1 protomer at an RMSD of 2.3Å

To evaluate the ability of AF2 to predict the tertiary structure of CAV1 protomers, we compared the structure of a single CAV1 protomer extracted from the experimentally determined structure (PDB-7SC0) with a model of the CAV1 monomer structure predicted by AF2 (AF-Q03135-F1, **Figure 1**). Several regions of CAV1 (residues 1-48 and 178) were predicted with low confidence. These regions overlap with the region of CAV1 that were not resolved in the experimental structure [11]. We thus focused our comparison on the well resolved/high confidence regions of the experimental and predicted structures. Overall, there was marked similarity between the predicted and experimental structures, with a few notable exceptions (**Figure 1**). First, the experimental protomer contains two α-helices (α3 and α4) separated by a short loop close to the boundary between the IMD and SR, whereas the predicted CAV1 AF2 monomer forms a continuous α-helix with a slight kink near residue P132 in the same region. The predicted structure also is missing a loop between α4 and α5. The absence of these loops gives rise to a difference in curvature between the two structures (**Supplementary Figure 1**). Furthermore, the last ten residues are predicted to form a coiled C-terminus in the AF2 monomer structure, in contrast to the experimentally observed β-strand geometry of the protomer (**Figure 1**). Nevertheless, considering that the AF2 model for CAV1 was predicted in the absence of an actual structure for CAV1 in the PDB, combined with the fact that the CAV1 monomer exists in nature within a higher order complex, the degree of similarity between the two structures is remarkable.

### AF2 predicts the overall quaternary structure of CAV1 oligomers

We next asked whether AF2 is capable of assembling CAV1 protomers into closed discs, and if so if it correctly predicts CAV1 exists as an 11-mer. To test this, we generated *n*-mers containing between 2 to 15 copies of CAV1 using AF2.2 (**Figure 2, Supplementary Figure 2**). The probability of forming a closed structure varied across n-mers (**Supplementary Figures 2 and 3**). Oligomers containing six or fewer CAV1s were unable to generate closed assemblies where the pin motif was properly positioned (**Figure 2, Supplementary Figure 2**). A closed assembly was observed for the 7-mer, but its structure was highly curved compared to the experimental Cav-1 structure, and the best prediction for the 8-mer failed to form a closed disc. However, the best fitting models of oligomers containing between 9 and 15 protomers were predicted to form closed discs with an architecture similar to the experimental CAV1 11-mer structure (**Figure 2, Supplementary Figure 2**). The top model predictions were highly reproducible for the case of the 11-mers (**Supplementary Figure 4**). The complexes formed by the 9-to 15-mers share several structural features. Similar to the experimental structure, the PM of each protomer locks neighboring CAV1 protomers in place along the outer rim of the complex. The predicted structures also exhibit extensive interactions between the hydrophobic residues of α-helices α2, α3, and α4. The last nine amino acids of the C-termini of each protomer form β-strands that assemble into a parallel β-barrel. This β-strand was not observed in the CAV1 monomer structure predicted by AF2 as measured from the Φ and Ψ angles at this region (**Figure 1F, Supplementary Figure 5**), suggesting it forms as a consequence of the oligomerization process.

**Figure 2.**
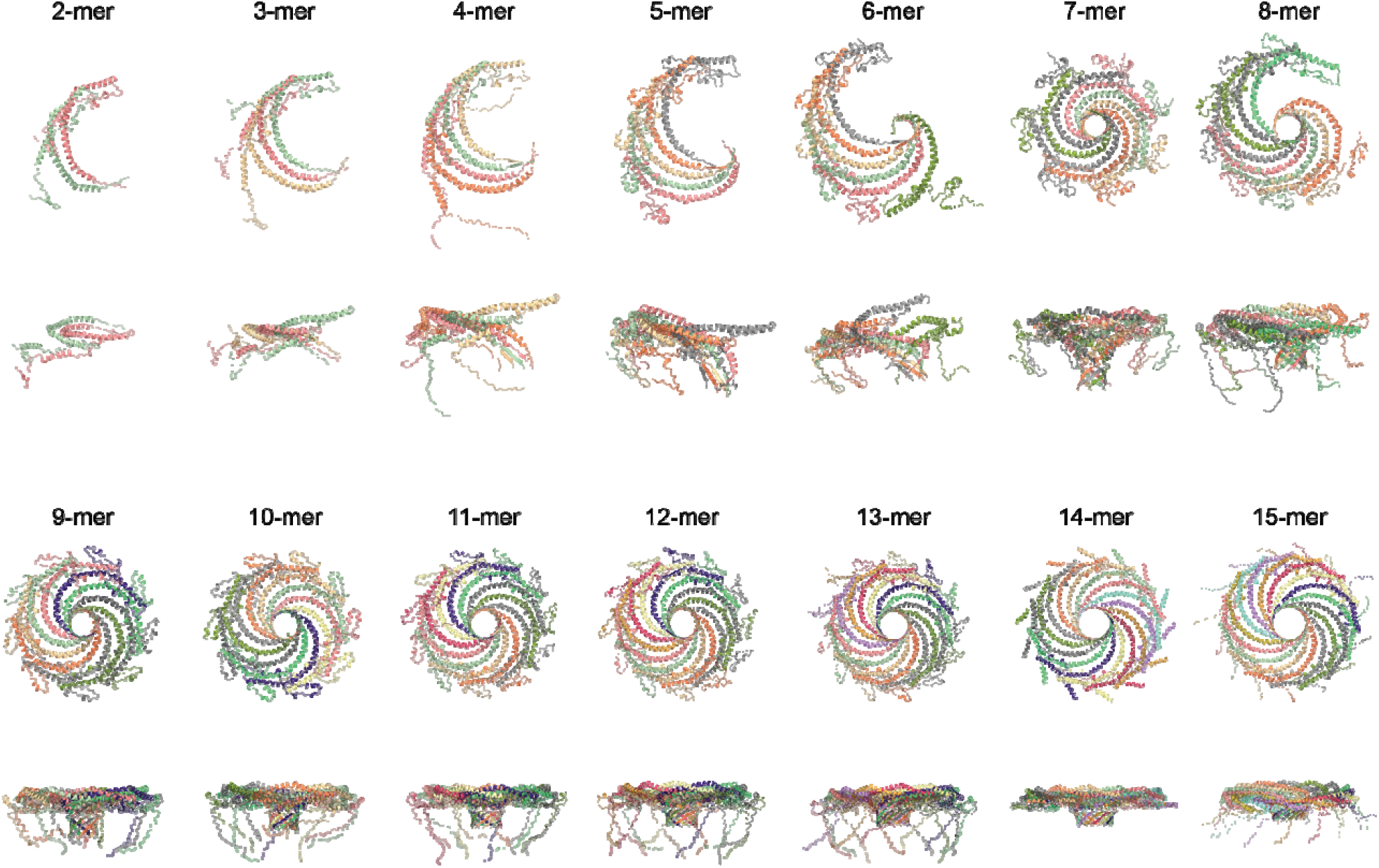
Best fit predictions by AF2.2 for CAV1 n-mers ranging in size from 2-15. *En face* (top) and side views (bottom) are shown for each case. Closed discs are predicted to form for n = 7 and n= 9-15. For the 2-to 13-mers, the predictions were based on full length alpha isoform (1-178) of human CAV1. Predictions for the 14- and 15-mers were based on the sequence of the beta isoform (32-178) of human CAV1. Each protomer is depicted in a different color. All structures are shown to scale.

We also compared the predictions of AF2.2 and AF2.1. Like AF2.2, AF2.1 predicted CAV1 can assemble into closed discs, but in this case only 7 or 8 CAV1 protomers were required. Furthermore, oligomers composed of 9 or more protomers exhibited significant clashes and unrealistic structures (**Figure 3**). Thus, AF2.2 was used for all subsequent analysis.

**Figure 3:**
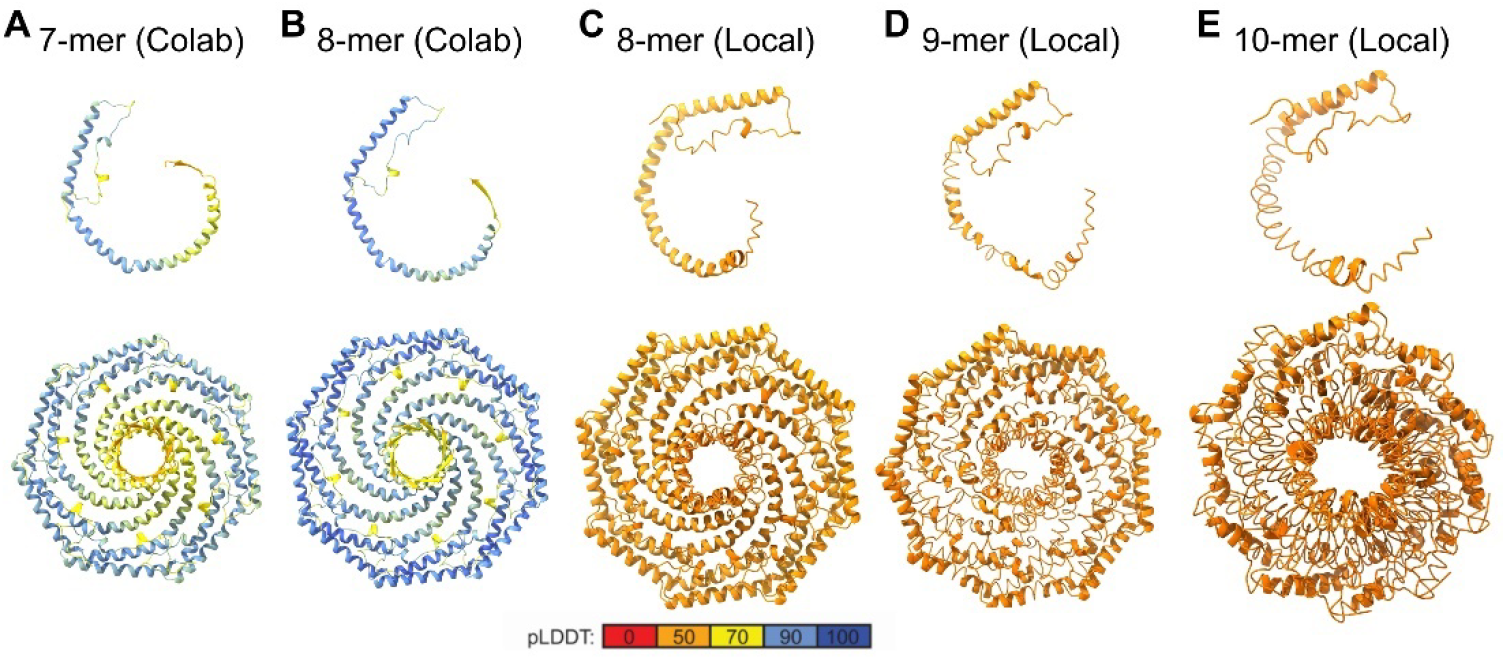
CAV1 oligomers predicted by AF2.1 become increasingly distorted as a function of increasing number of protomers in the complex. Predicted structures for CAV1 oligomers generated with the Colab version of AF2.1 (modeled as 32-178) and a local installation of AF2.1 (modeled as 1-178). (**A** and **B**) 7- and 8-mer CAV1 generated with the Colab version, (**C-E**) 8-, 9-, and 10-mer generated with a local installation of AF2.1. A protomer belonging to each oligomer is shown in the top row and the whole complex viewed en face is shown in the bottom row. Only residues 49-178 are shown for clarity. All structures are shown to scale.

### Conserved critical interactions at protomer – protomer interfaces stabilize CAV1 oligomers with 10 to 12 copies

We next investigated the protomer-protomer interactions that underlie the formation of the CAV1 disc structure repeatedly observed among the AF2-predicted structures in more detail (**Figure 4**). For these studies we chose to analyze 10-, 11-, and 12-meric models generated by AF2 due to their similarity in size to the 11-meric experimental CAV1 structures (**Figure 4A-L**). Alignment of the AF2-predicted protomers showed a similar tertiary structure that mainly differed in terms of the curvature of the α-helical regions, as well as shifts in the positioning of the PM region that locks the neighboring protomer in place at their turning point (**Figure 4M,O**). The curvature differences result from the presence of several turns in the SR in the experimental CAV1 protomer that are not observed in the predicted protomer structure (**Figure 5, Supplementary Figure 1**), similar to the case of the AF2 CAV1 monomer. The differences between the curvature of the SR explains how the predicted and experimental CAV1 structures can assemble differently at the quaternary level. Several additional differences between the experimental and predicted structures were noted. In contrast to the flat membrane facing surface of the cryoEM structure, the membrane facing surface of the predicted complexes bows in toward the center of the complex (**Figure 4C, F, I, L**). The β-barrels of the AF2 models are positioned above the outer rim of the complex (**Figure 4B, E, H, K**). Finally, the overall diameter of the predicted complexes is also slightly smaller than that of the experimental complex (133 Å for the predicted 11-mer versus 140 Å for the cryoEM structure).

**Figure 4:**
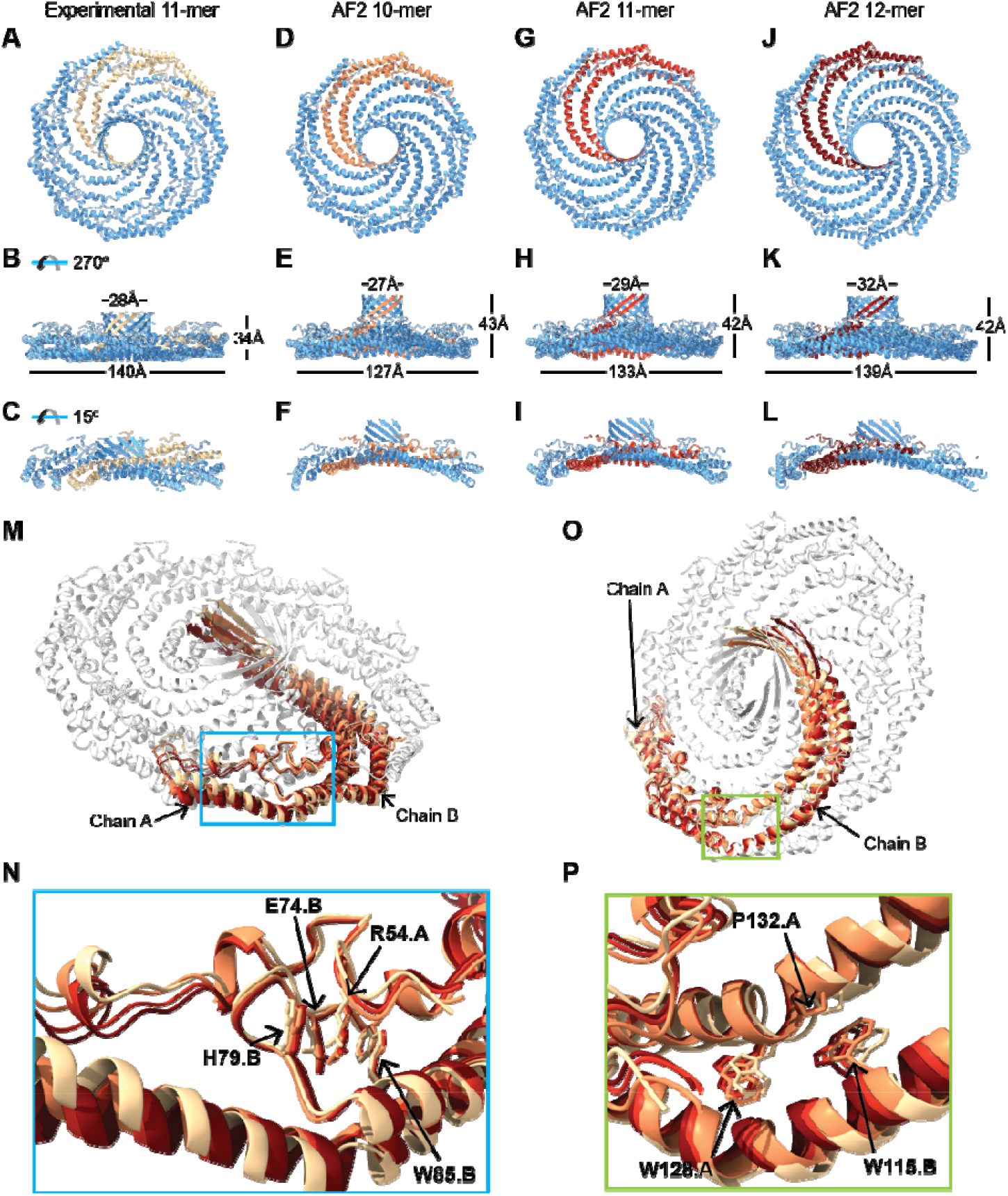
Packing and interactions of neighboring protomers in predicted CAV1 complexes are similar to their counterparts in the cryo-EM based structure. **(A-L)** Comparison of *en face* and side views of secondary structure model of cryo-EM based CAV1 11-mer complex with structures of CAV1 10-, 11- and 12-mers predicted by AF2.2. Dimensions of complexes are labeled in B, E, H and K. Cut though views of each model are shown in panels C, F, I, and L. Residues 1 to 48 and 178 were hidden from the AF2.2 predicted structures to make them more comparable to the cryo-EM structure. Two neighboring protomers from each model are highlighted in gradually warming colors. All structures are shown to scale. **(M and O)** Overlay of two neighboring dimers extracted from each of the models shown in A-H. Protomers from the cryoEM structure are shown in beige or white. **(N and P)** Close up of boxed regions from panels M and O showing zoomed views of the key residues in the pin motif (N) and in surrounding P132 (P).

**Figure 5.**
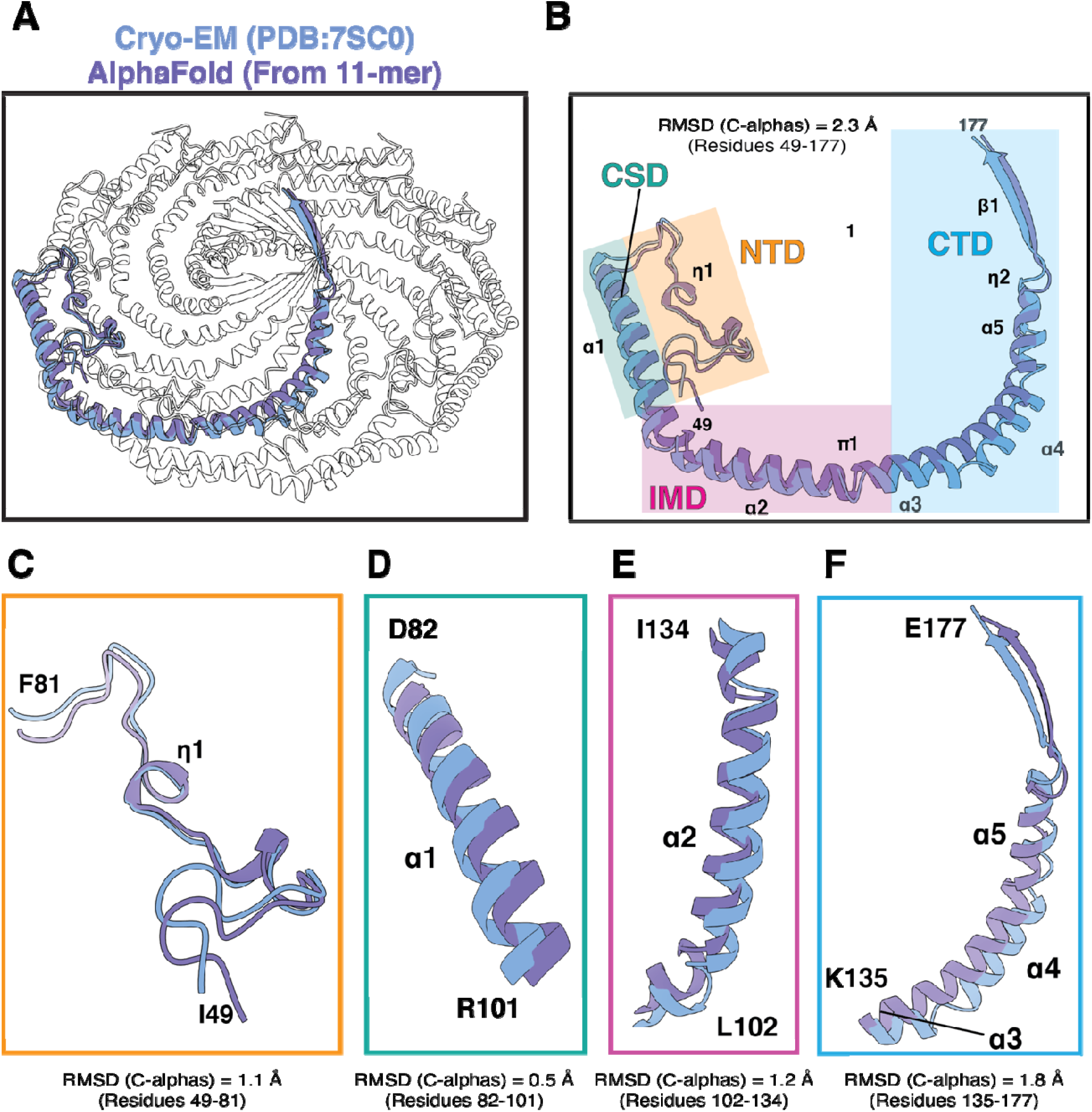
Comparison of the CAV1 protomers from the cryo-EM structure and 11-meric complex of CAV1 predicted by AF2.2. (**A)** The CAV1 8S complex (PDB 7SC0) is shown with a single protomer colored blue and overlayed with an aligned AF2.2 protomer taken from the AF2.2 predicted CAV1 11-mer in purple. (**B**) Overlay of a single protomer from the cryo-EM structure and the AlphaFold2 CAV1 protomer. The secondary structure assignments for the cryo-EM structure are labeled. The coordinates were aligned by their alpha carbons and their RMSD values are shown. The first 48 residues were not included in the alignments. Each region is highlighted by a different color. Orange, residues 49-81 of the N-terminal domain (NTD), teal, SD; pink, intramembrane domain (IMD), and blue, residues 135-177 of the C-terminal domain (CTD). (**C-F**) Each of the indicated regions was aligned individually and RMSD values calculated. Experimental structures are shown in blue and predicted structures are shown in purple.

We next focused on the neighborhood of two residues important for CAV1 structural integrity and function. The first one is R54 on the N-terminal α-helix forming the PM domain (**Figure 4N**). In the experimental structure, this arginine is sandwiched between the residues W85 and H79 of the neighboring protomer through π-π stacking interactions and forms additional electrostatic interactions with E74. Mutations of R54 were shown to abolish the ability of CAV1 to form disc-shaped assemblies in vivo, suggesting a key role for the stabilization of protomer – protomer interactions [11]. The other residue is P132, located close to the transition between the IMD and SR (**Figure 4P**). A P132L mutation at this site results in disruption of CAV1 structure in both monomeric and oligomeric environments [16, 17].

Comparisons of the three AF2 models and experimental structure showed that the vicinities of R54 and P132 are almost identical to the WT structure with all key interactions conserved. All three binding partners of R54 had nearly identical configurations among the three oligomeric states investigated (**Figure 4N**). In the case of P132, the relative coordinates of α-helix α3 shifted due to the changes in the curvature at the turning point of P110 connecting the IMD α-helices α2 and α3. However, hydrophobic interactions of P132 with W115 of the neighboring protomer were conserved in all three cases (**Figure 4P**). Further, the residue – residue interactions between neighboring protomers was nearly identical for all predicted CAV1 oligomers formed by 9 to 15 protomers, which was also consistent with the contacts calculated for the experimental 11-meric CAV1 structure with minor differences at the PM region (**Supplementary Figure 6**).

### CAV1 oligomers of different sizes show similar energetics at α-helical regions but vary in their ability to form stable β-barrels

The finding that AF2 predicts between 9 and 15 CAV1 protomers can assemble into oligomeric complexes with a conserved architecture raises the question of whether these *n*-mers differ in stability, and, if so, what regions of the protein contribute most importantly to these differences. To this end, we calculated the per-residue energies of each complex with Rosetta [18]. To account for interactions of CAV1 with membranes, we incorporated an implicit membrane model [19, 20] based on the distribution of residues in contact with detergent in the experimental structure [11]. The membrane-embedded residues consist of the helical domains of CAV1 whereas the β-barrel is predicted to be completely solvent-exposed. Because the first 48 residues of CAV1 are not observed in the experimental structure and are also predicted with low confidence by AF2, we confined our analyses to the CAV1 residues 49-178.

Both the experimental 11-meric CAV1 structure and the AF2 models had favorable energies overall, indicated by blue color throughout the per-residue energy distributions (**Supplementary Figure 7)**. A closer inspection of the individual domains showed that the PM had the largest number of residues with high energy, followed by the β-barrel and α4 as indicated by red color. Increasing oligomer size had a slight but noticeable effect on the stabilities of the helices α1 and α2, whereby these domains had lower energies in larger assemblies such as the 14- and 15-meric structures. The most striking difference was observed at the β-barrel region. Here, larger assemblies with thirteen to 15 protomers showed higher calculated scores suggesting issues with the stability of their β-barrels (**Supplementary Figure 8**). Further, the experimental CAV1 structure had lower energies in the β-barrel region compared to the AF2-generated models including the 11-meric AF2 model (**Figure 6**).

**Figure 6:**
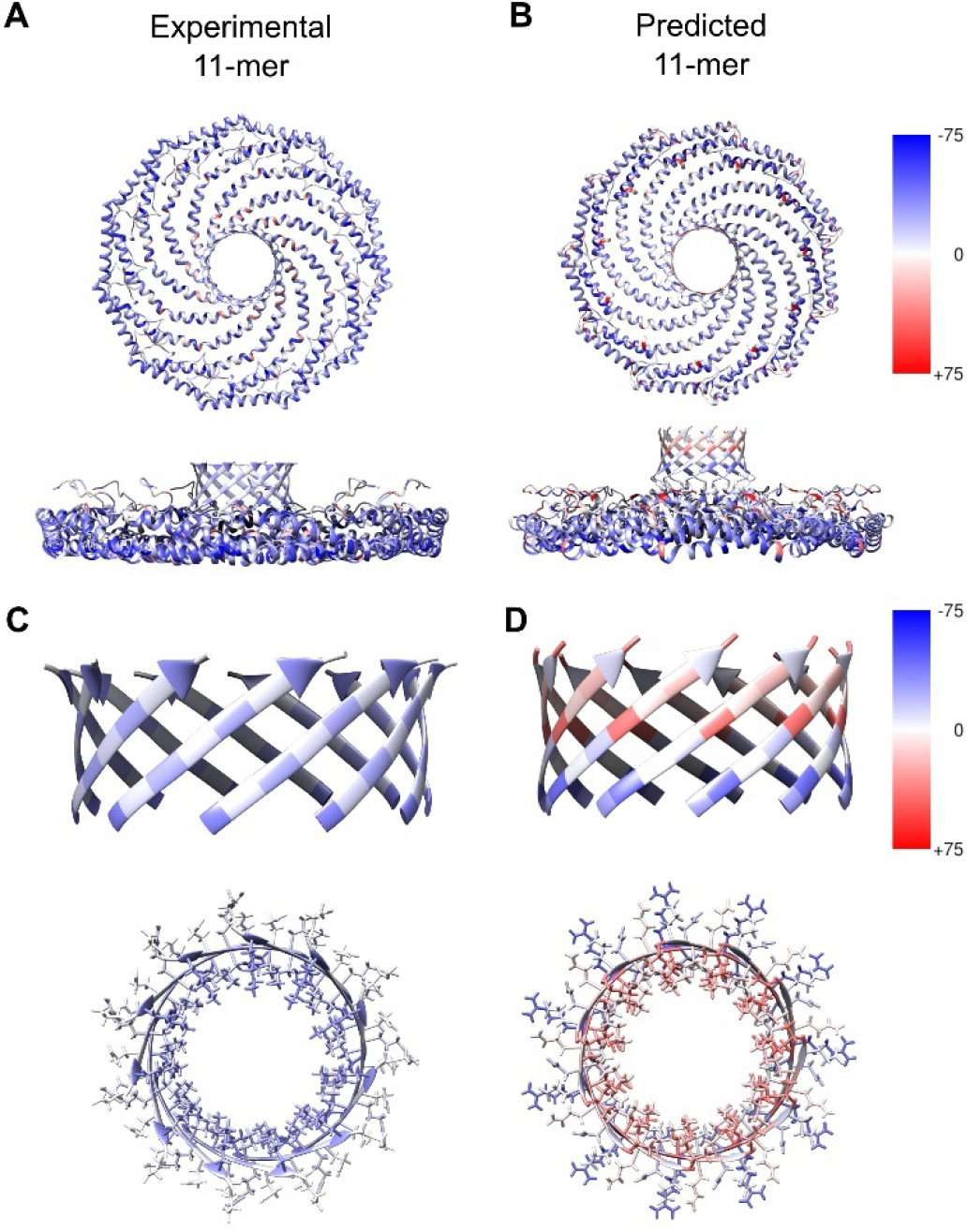
Comparison of per-residue energies for the experimental and AF2-generated 11-meric CAV1 structures. (**A, B**) Per-residue energies mapped onto the (A) experimental 11-meric CAV1 structure versus the (B) AF2-predicted structures the 11-meric complex. (**C, D**) Close up view of energetics of CAV1 β-barrel residues for experimental and predicted structures. Red indicates unfavorable energies and blue indicates favorable energies. All scores are in Rosetta Energy Units (REU). Structures in A and B are shown to scale.

### Unusual design principles underlie CAV1’s 11-stranded parallel β-barrel

The central parallel β-barrel of CAV1 is formed by alternating hydrophobic residues lining the cavity and hydrophilic residues facing outward. One exception to this pattern is an inwardly facing lysine residue at the position 176, a highly conserved residue amongst CAV1s. An evolutionary analysis of 315 CAV1 proteins suggests the propensity to form β-barrels with a similar pattern of residues is conserved across different species, highlighting the importance of the alternating hydrophobicity motif for the structure and function of the protein (**Supplementary Figure 9**). In all AF2 models, the residues facing the barrel pore had higher energies compared to the outer facing residues (**Supplementary Figure 8, bottom views**). This reflects the tight packing of the inner barrel residues, which are positioned at the same level for each strand as a consequence of its shear number of 22. Another interesting finding was the visible distortions observed for the barrels larger than twelve strands, with accompanying high calculated energies (**Supplementary Figure 8**). The experimental CAV1 β-barrel had a more uniform distribution of scores for the inside and outside facing residues with a lower overall energy compared to the predicted models. The experimental β-barrel also had a lower overall energy compared to the predicted structures, which may partly be attributed to its larger barrel diameter (∼26.5 Å in predicted 11-mer vs. ∼30 Å in experimental) and more favorable alignment of the pore-facing residue rotamers compared to the predicted structures.

Lastly, we compared the energetics of the CAV1 barrel with two other parallel β-barrels to assess the effects of barrel size on their stability (**Figure 7, Table 1**). Most parallel β-barrels in nature belong to the TIM-barrel family whose members consist of eight β-strands that fold typically into a barrel with C4 symmetry [21] (**Figure 7**). Another example was recruited from a soluble phage protein (*Bacillus cereus*) whose C-termini folds into a central 9-meric parallel β-barrel (2AO9) (**Figure 7**). In comparison to the 9-meric barrel, a representative 8-meric TIM barrel (5BVL) shows a more stable energy profile. The terminal residues of both the selected 8-mer and 9-mer barrels are asymmetric, likely to help prevent clashes caused by the proximity of pore-facing residues in these tight assemblies. In contrast, the CAV1 barrel is perfectly symmetric, suggesting its larger radius allows less strain between the residues pointing into the barrel (**Figure 7**). Although the inner lining of the TIM barrel example and CAV1 had mostly polar residues deeper in the pore, the 9-meric β-barrel had no clear preference. The 8-meric structure had a tight pore caused by the tightly packed hydrophobic residues at the center of the pore and arginine residues at its C-terminus. The pore size was larger for the 9- and 11-meric β-barrels, as expected. Comparison of the 9-meric β-barrel with the AF2-predicted 9-meric CAV1 structure showed marked similarities at the barrel region potentially due to the constrained nature of a parallel β-barrel, but the geometry of other domains had marked differences (**Supplementary Figure 10**).

**Figure 7:**
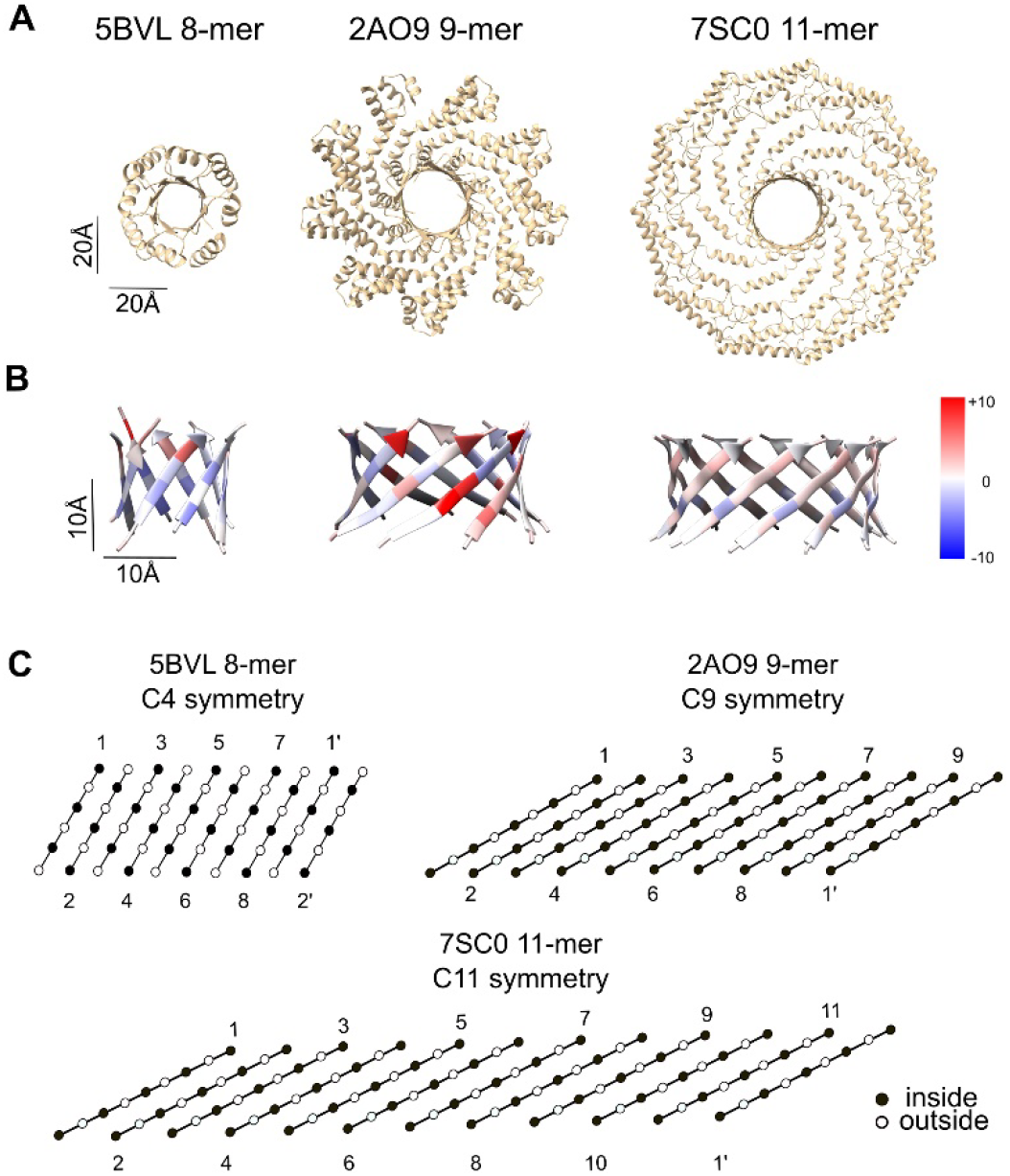
Comparison of structural features of CAV1 beta barrel with other parallel beta barrels. (**A**) The full structures of the assemblies from which the analyzed parallel β-barrels were taken viewed in the direction of the barrel pore. All structures are shown to scale. (**B**) Structures and the per-residue energy breakdowns of three parallel β-barrels calculated with Rosetta. Left panel, a representative TIM barrel with C4 symmetry (PDB 5BVL); middle panel, a phage protein from Bacillus cereus with C9 symmetry (PDB 2AO9); right panel, experimentally determined CAV1 β-barrel with C11 symmetry (PDB 7SC0). Red indicates positive scores (destabilizing) and blue indicates negative scores (stabilizing). (**C**) Amino acid alignments of the 8-, 9-, and 11-meric β-barrel structures. Black circles indicate pore-facing residues and white circles indicate residues facing away from the pore. Individual strands are labeled. Primes denote the same strand as the starting strand and are included to show the relative levels of the first and last strands in the barrel.

**Table 1.** Characteristics of three different parallel β-barrels analyzed. Number of strands (n), shear number (S), register shift (t), barrel radius (R) measured between the farthest α-carbons, and the angle between an individual β-strand and the major helix axis (θ).

| **PDB ID** | **n** | **S** | **t** | **R (Å)** | **$\theta$ (°)** |
| --- | --- | --- | --- | --- | --- |
| 5BVL | 8 | 8 | 1 | 8.5 | 37 |
| 2AO9 | 9 | 18 | 2 | 14 | 48 |
| 7SC0 | 11 | 22 | 2 | 15 | 53 |

## Discussion

Even for an isolated monomer, AF2 was able to predict the tertiary structure of CAV1. This is surprising as the biophysical context was absent, i.e. other protomers to form the assembly, and the membrane environment that is likely to play a significant role in CAV1 folding considering its role as a membrane-embedded curvature-inducing protein. It is particularly astounding that AF2 predicts this fold unambiguously, also reflected by the uniformity of the monomeric predictions in our calculations, as one would intuitively expect a large ensemble of possible conformations with rather similar energy. Apparently, local conformational preference is sufficient to determine the structure of the protomer even in isolation. Interestingly, and consistent with the absence of hydrogen bonding partners to form the β-barrel, AF2 folded the C-terminal residues into a coil.

The 11-meric CAV1 structure predicted by AF2 was consistent with the general scaffold of the experimental CAV1 structure. Interestingly, while some detailed aspects of the structure were predicted consistently at atomic-detail accuracy, such as helix-helix interactions, other overarching features were predicted in an inconsistent manner, such as the disc-shaped symmetric arrangement and the curvature and length of the α-helical regions. We argue that this hints at an important metric of AF2 multimer: It is achieving a high local accuracy driven by the hundreds of thousands of protein structures it has been trained with while sometimes struggling with global aspects of novel arrangements of these local features [10].

Although the experimental structure of CAV1 suggests it prefers to exist as an 11-mer, AF2 predicts that the protein has the capacity to assemble into oligomers containing between 7 and 15 protomers. This range of values overlaps remarkably well with early estimates of the oligomeric state of CAV1 [22-24]. Comparison of the predicted models reveals how the complex could potentially accommodate different copy numbers of CAV1. Many interactions between the domains of neighboring protomers are independent of the number of protomers forming the oligomers in the range of 9-15 protomers. All oligomers successfully formed the PM that locks neighboring CAV1 protomers. Hydrophobic interactions between the neighboring IMDs were also conserved, although the angles at the turning points varied among the different protomer numbers, which caused curvature differences at the SR. These finding suggest that other oligomeric forms of CAV1 could exist in nature and may be selected for under different physiological conditions.

Formation of a closed and well-defined β-barrel likely helps to define the oligomeric state of the CAV1 complex. Of the predicted 7 to 15-meric complexes, all but 8-meric CAV1 are capable of doing so. Oligomers containing 10-12 protomers had the highest probability of forming discs, and the β-barrel was most stable with 9 to 12 copies. Further, the experimental 11-meric CAV1 structure had a more stable β-barrel compared to the AF2-generated models. These results may indicate that while interactions at the PM may be important for the formation of CAV1 oligomers, the overall size and stability of the complex is heavily dependent on the stability of the β-barrel.

Our findings also uncover design features that enable CAV1 to assemble into such large parallel β-barrels. By their nature, β-barrels are constrained by the hydrogen bonding requirements for the formation of the barrel shape [25]. The majority of the parallel β-barrels found in the PDB are TIM barrels, i.e. α_8_β_8_-barrels, formed by eight β-strands and eight α-helices. Interestingly, α_8_β_8_-barrels have (only) a four-fold symmetry, as sidechains from uneven β-strands point inward while sidechains from even β-strands point outward, and vice versa. The highest symmetric β-barrel in the PDB prior to CAV1 is a β-barrel consisting of nine β-strands that is formed by the C-terminal regions of a protein from Bacillus cereus (PDB ID: 2AO9, **Supplementary Figure 10**). Comparison of these β-barrels with the experimental CAV1 structure showed that the sidechains of the former structures break the symmetry of the system by shifting of the pore-facing side chains to adapt to the tight packing environment inside the pore, whereas the CAV1 β-barrel showed full symmetry likely due to its larger pore size (∼19 Å, ∼24 Å, and ∼28 Å). These findings, when taken together with the AF2 predictions for CAV1 oligomers with different numbers of protomers, suggest that parallel β-barrel sizes formed by 10 to 12 β-strands are best tolerated when the same residue of each β-strand is pointing inside simultaneously.

Overall, these results show the effectiveness of AF2 as a tool to predict the structures of novel overall oligomeric protein architectures even if the protomers have no close homologues in the PDB, despite some limitations regarding the prediction of secondary, tertiary, and quaternary structure. Extension of analyses such as ours to all the putative members of the caveolin family may not only enable us to better understand the conserved motifs among this family, but also provide a useful screening tool to distinguish sequences that are annotated as caveolins based on their tendencies to form disc-shaped structures similar to observed for human CAV1. Importantly, while the experimental structures of CAV1 displays an 11-mer, other arrangements of open and closed discs are likely to be biologically relevant and might be similar to the AF2 proposed models. Such structures could, for example, potentially correspond to intermediate complexes formed as the newly synthesized protein first assembles into oligomers.

Finally, improvements to the oligomeric prediction capabilities of AF2 will drastically improve our ability to predict protein – protein complex structures in the absence of experimental data regarding the interaction sites of such multimeric assemblies.

## Methods

### AF2 calculations

Prediction runs were executed either using AlphaFold v2.2.0 with default settings via a Colab notebook named “alphafold21_predict_colab” provided by ChimeraX daily builds version (ChimeraX 1.4.0) or using AlphaFoldv2.1 via another colab notebook named “AlphaFold2_advanced (AlphaFold2_advanced.ipynb - Colaboratory (google.com)” with default settings. Full-length human CAV1 was used for AlphaFold v2.2.0 predictions for structures containing two to thirteen protomers. Due to memory limitations, the sequence of human CAV1-beta-isoform (residue 32-178) was used for the predictions of 14-mer and 15-meric CAV1. For AlphaFold v2.1, predictions for 8-mer complexes were based on the sequence of human CAV1-beta-isoform.

The figures were analyzed, rendered and exported with either ChimeraX 1.4.0 (daily builds version) or ChimeraX1.3. RMSD values were calculated using ChimeraX1.3. Chain A from the model was used as a reference for all structure matching.

### Rosetta cryo-EM relax calculations

The experimental CAV1 structure was relaxed with the Rosetta cryo-EM relax protocol [26]. A single protomer was relaxed with 11-fold symmetry based on symmetry files generated using the 11-meric experimental CAV1 structure. The *ref2015_cart* score function with residue weights adjusted for cryo-EM calculations was used to score the structures. An *elec_dens_fast* value of 50 was selected as the restraint weight for the relax calculations based on the resolution of the CAV1 structure. A total of 500 structures were generated and the lowest-scoring structure was used for the analyses based on lowest total score.

### Per-residue energy calculations

The per-residue calculations were run with Rosetta *per_residue_energies* application [18]. For each oligomeric assembly, membrane coordinates were calculated using the Positioning Proteins in Membranes (PPM) server [27]. The per-residue energies of the aligned protein structures were calculated with the *mpframework_smooth_fa_2012* score function [19, 20]. For the comparison of the three β-barrels from the PDB IDs 5BVL, 2AO9, and 7SC0, the soluble membrane score function *ref2015* was used to calculate the per-residue energies since these structures are not embedded in membrane [28]. The resulting energies from the per-residue energy calculations were used to color each oligomeric assembly using the Define Attribute module of UCSF Chimera. A three-color coloring scheme from blue to red color was used to represent the negative and positive per-residue scores respectively.

### Residue contact map predictions

The residue – residue contact map predictions were made and plotted with the Bio.PDB module of the BioPython software [29]. Two neighboring protomers of each oligomeric structure were used for the contact map analyses. The α-carbon coordinates were calculated for every residue in the system and the pairwise distances were calculated for each residue. Of the calculated distances, each pair within 12 Å of each other was considered as a contact, and only the pairs of residues in contact were plotted.

## Acknowledgements

We thank Erkan Karakas for helpful discussions.

