## Supplementary material for "Template-free prediction of a new monotopic membrane protein fold and oligomeric assembly by Alphafold2"

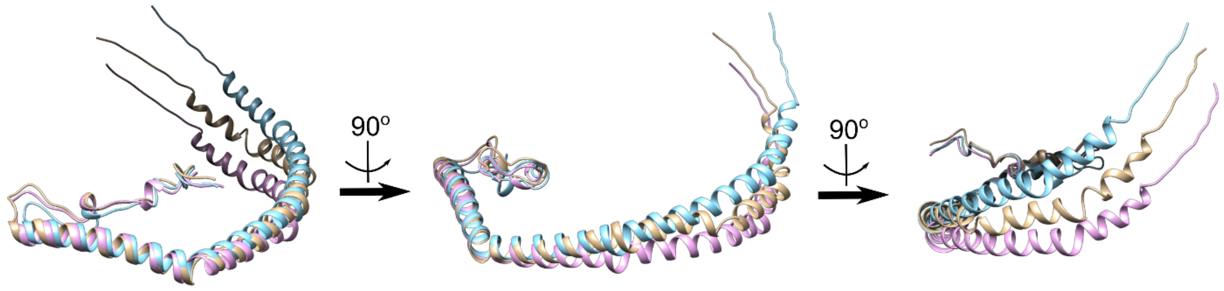


**Supplementary Figure 1**: **Comparison of experimental and predicted CAV1 protomer and monomer structures.** Alignment of the experimental 11-meric CAV1 protomer (beige), CAV1 AF2 monomer (blue), and CAV1 AF2 protomer from an 11-meric prediction (pink) using helix α1 as the anchor point viewed from three different angles.

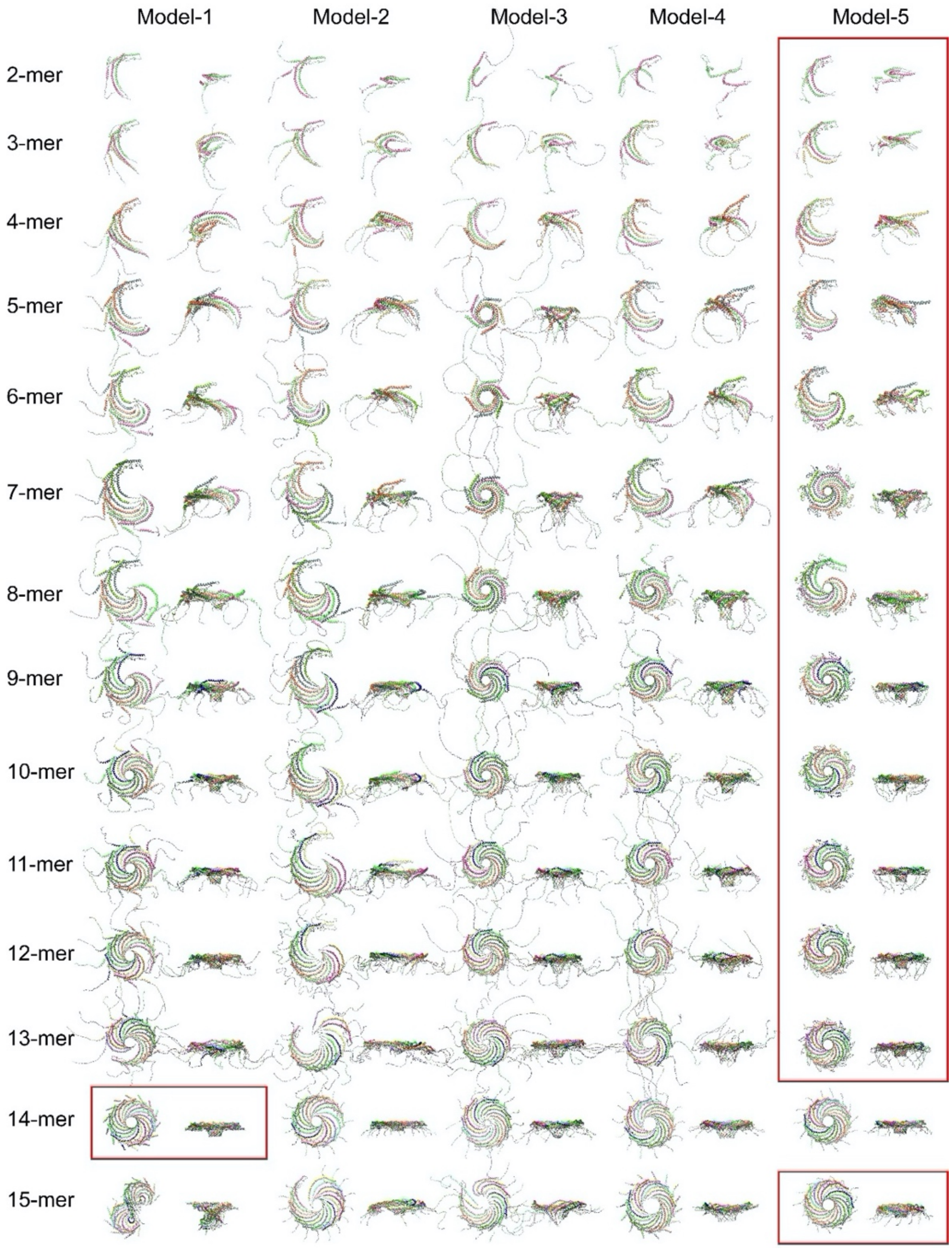


**Supplementary Figure 2: Top and side views of CAV1 homo-oligomers predicted by AF2.2.** Five models are show for each prediction. The best model from each prediction is outlined by a red box. For the 2- to 13-mers, the predictions were based on full length alpha isoform (1-178) of human CAV1. Predictions for the 14 and 15-mers were based on the sequence of the beta isoform (32-178) of human CAV1. Each protomer is indicated using a different color within a given complex.

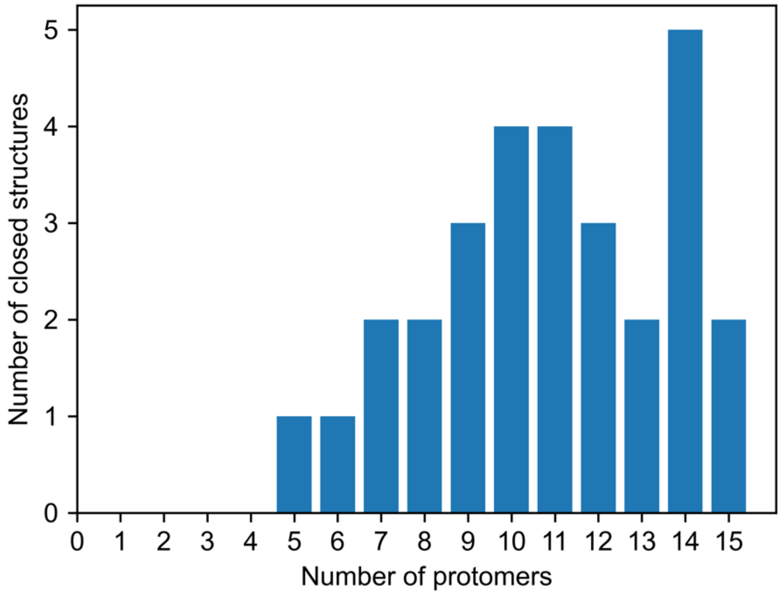


**Supplementary Figure 3**: **The probability of forming a closed structure varies across n-mers.** Distribution of the predicted number of closed structures (out of the top 5 predictions) as a function of the number of input CAV1 protomers. A closed structure is defined as an oligomer in which all protomers folded into a closed disc-shaped assembly.

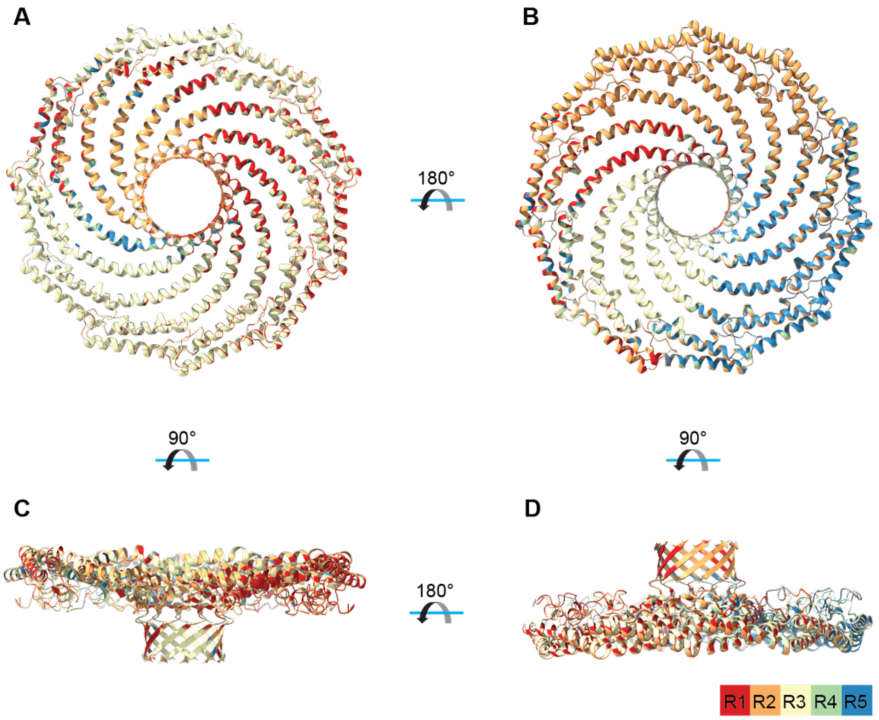


**E** RMSD between repeated predictions

| | R2 vs. R1 | R3 vs. R1 | R4 vs. R1 | R5 vs. R1 |
|---|---|---|---|---|
| **49-177** | *0.034* | *0.032* | *0.037* | *0.038* |

**Supplementary Figure 4: AF2.2 predictions for CAV1 11-mers are highly reproducible. (A-D)** Overlay of five independent predictions of best fitting AF2.2 models of the CAV1 11-mer (predicted using residues 49-177)**.** The best fitting model for each repeat (R1-R5) is shown in a different color, as indicated in the legend. Four different views of the complex are shown. **(E)** Table of RMSD values compared across independent repeat predictions (repeats 1 through 5). Comparisons were all made with respect to repeat 1 (R1).

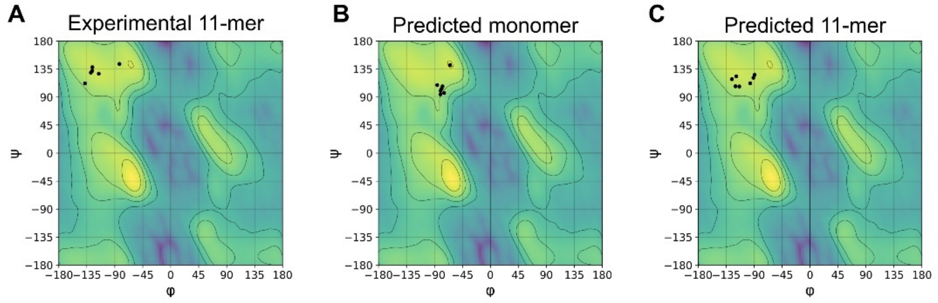


**Supplementary Figure 5**: **Ramachandran plots for residues within the β-strand region of the experimental CAV1 protomer structure.** (**A**) Protomer from the experimental CAV1 11-mer structure. (**B**) AF2-predicted CAV1 monomer. (**C**) Protomer from the AF2-predicted CAV1 11-mer.

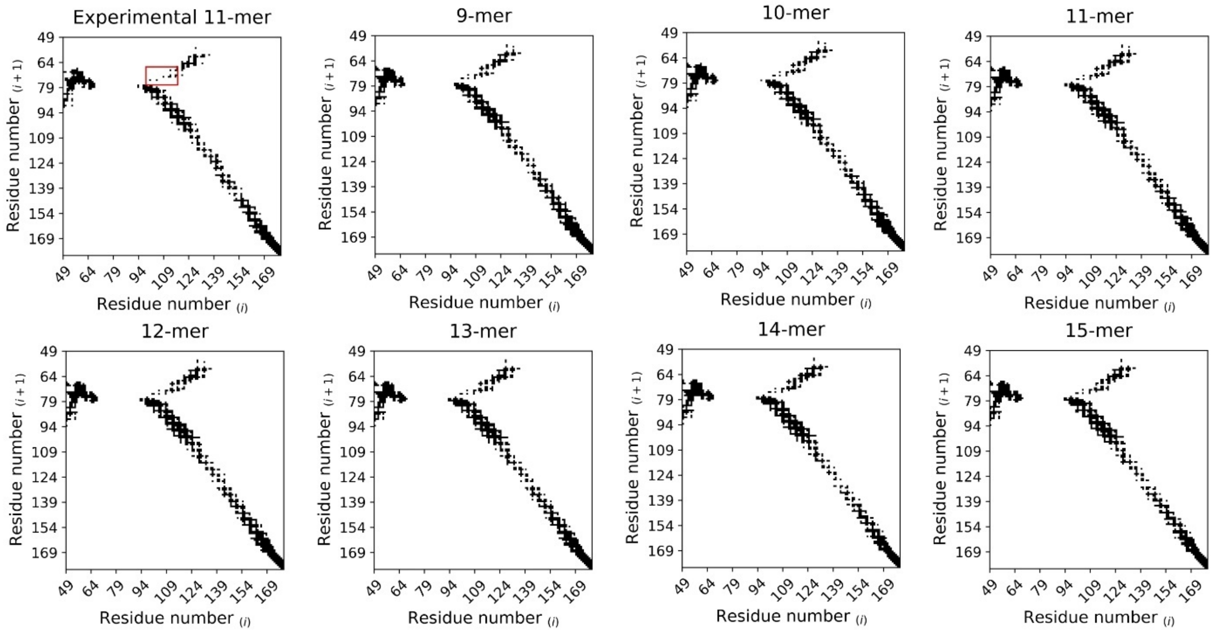


**Supplementary Figure 6: Contact maps calculated for different oligomeric assemblies of CAV1 including the 11-meric experimental CAV1 structure and the AF2.2-predicted oligomers consisting of 9 to 15 protomers.** A contact is considered as the proximity of any two residues within 12 Å of each other. The x-axis stands for the residue number at any given protomer, and the y-axis stands for the residue numbering at the neighboring protomer in the counter-clockwise direction. The region that showed increased contacts in AF2 models compared to the experimental structure is highlighted with a red box.

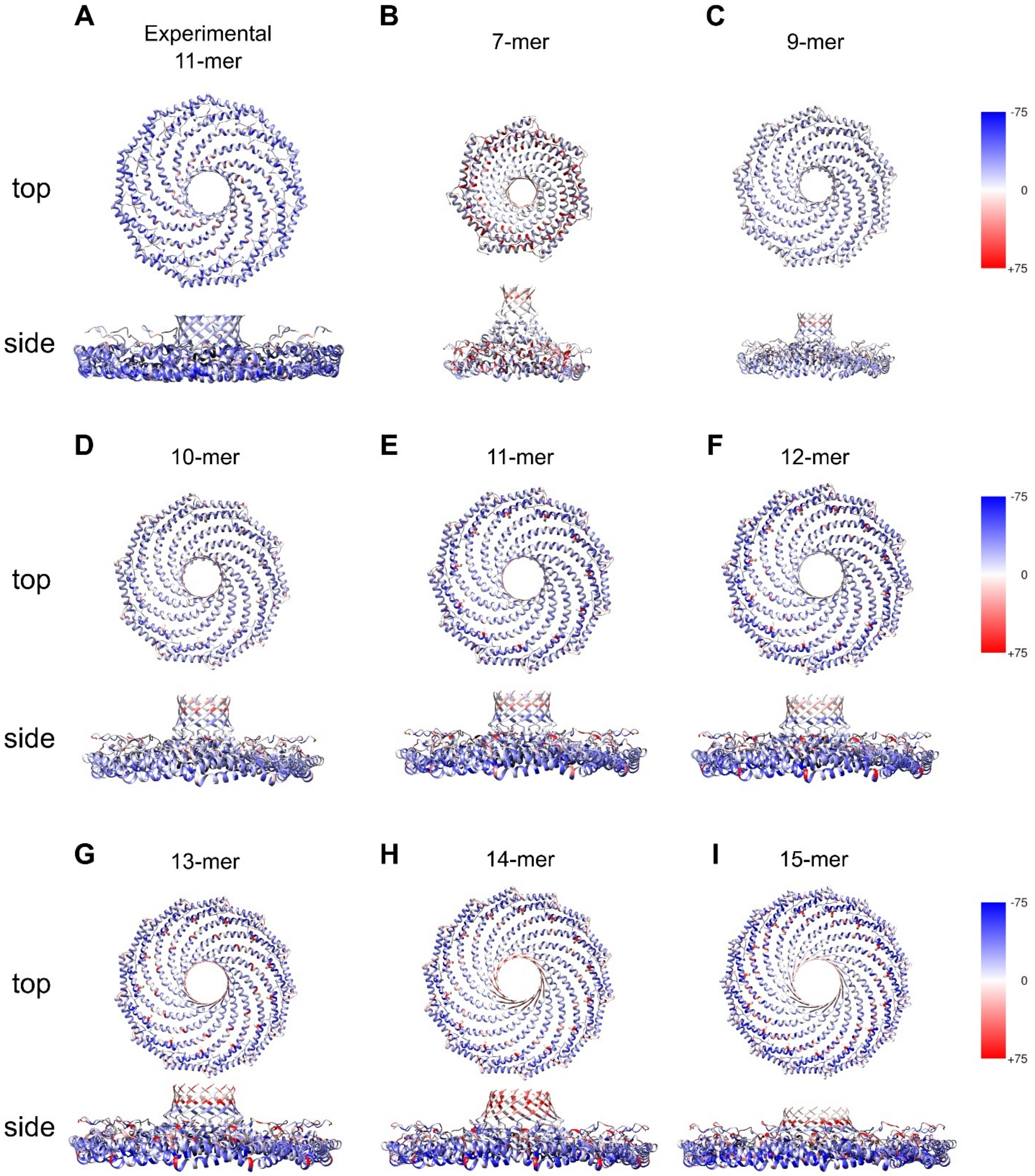


**Supplementary Figure 7: Per-residue energies calculated for the experimental 11-meric CAV1 structure and the AF2.2-generated CAV1 structures consisting of seven to 15 protomers.** Red color indicates unfavorable energies and blue color indicates favorable energies. All scores are in Rosetta Energy Units (REU). All structures are shown at the same scale.

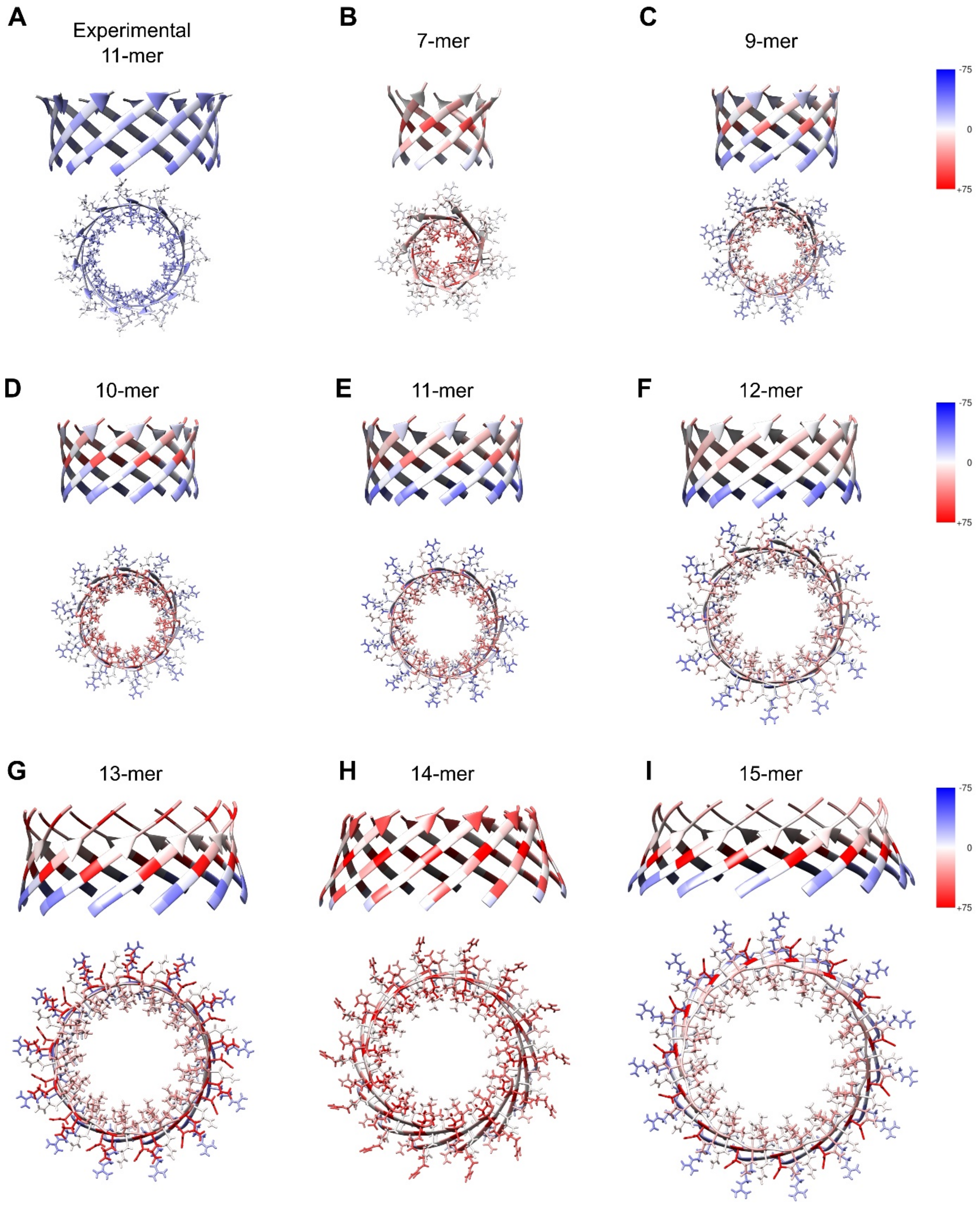


**Supplementary Figure 8: Per-residue energies for CAV1 β-barrels.** Shown are per-residue energies calculated using Rosetta for experimental 11-meric CAV1 structure and the AF2-predicted CAV1 oligomers from 7 to 15 protomers. Red color indicates unfavorable energies and blue color indicates favorable energies. All scores are in Rosetta Energy Units (REU). All structures are shown at the same scale.

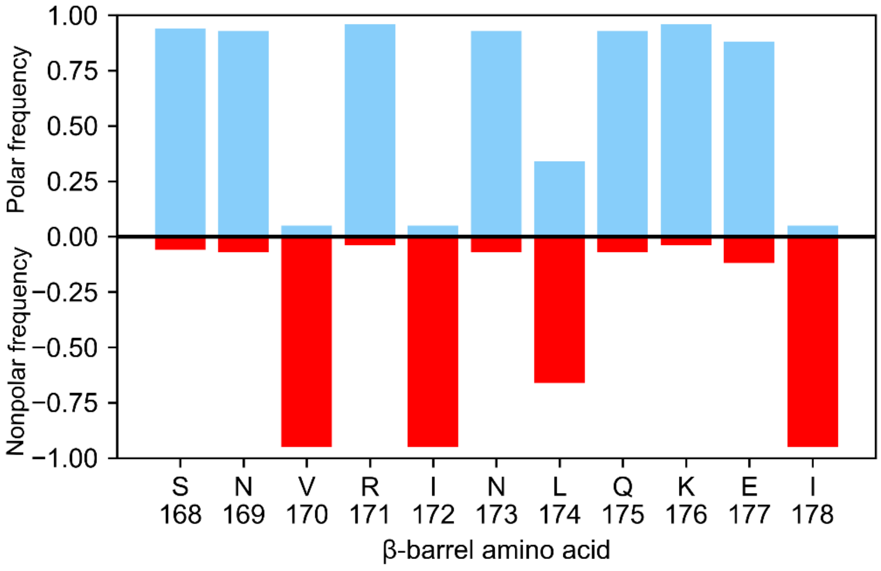


**Supplementary Figure 9: The presence of alternating polar and non-polar residues, together with a lysine near the C-terminus is a conserved motif found in the β-barrel region of CAV1 across multiple species.** Polar amino acid frequencies were calculated for the sequences of 315 CAV1 from different species. The x-axis shows the corresponding sequence from human CAV1 (residues 168-178), and the y-axis reports the probability of having a polar residue (blue) or nonpolar residue (red) at the corresponding position among different CAV1 sequences.

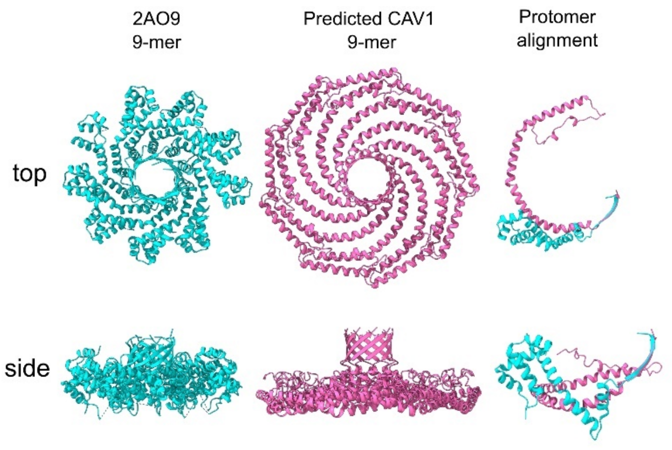


**Supplementary Figure 10**: **Comparison of the 9-meric experimental 2AO9 structure (cyan) and the AF2-predicted 9-meric CAV1 structure (pink) viewed from top (top row) and side (bottom row).** Alignment of a single protomer from each structure based on β-barrel coordinates is shown in the right panel. All structures are shown at the same scale.
